# Optimized cross-linking mass spectrometry for in situ interaction proteomics

**DOI:** 10.1101/393892

**Authors:** Zheng Ser, Paolo Cifani, Alex Kentsis

## Abstract

Recent development of mass spectrometer cleavable protein cross-linkers and algorithms for their spectral identification now permits large-scale cross-linking mass spectrometry (XL-MS). Here, we optimized the use of cleavable disuccinimidyl sulfoxide (DSSO) cross-linker for labeling native protein complexes in live human cells. We applied a generalized linear mixture model to calibrate cross-link peptide-spectra matching (CSM) scores to control the sensitivity and specificity of large-scale XL-MS. Using specific CSM score thresholds to control the false discovery rate, we found that higher-energy collisional dissociation (HCD) and electron transfer dissociation (ETD) can both be effective for large-scale XL-MS protein interaction mapping. We found that the density and coverage of protein-protein interaction maps can be significantly improved through the use of multiple proteases. In addition, the use of sample-specific search databases can be used to improve the specificity of cross-linked peptide spectral matching. Application of this approach to human chromatin labeled in live cells recapitulated known and revealed new protein interactions of nucleosomes and other chromatin-associated complexes in situ. This optimized approach for mapping native protein interactions should be useful for a wide range of biological problems.

## Introduction

Protein-protein interactions mediate cellular functions including signaling, metabolism and cell differentiation. Disruption of protein-protein interactions can contribute to disease and defining how these interactions change across the proteome can provide important insights into pathogenic mechanisms and ultimately improved therapies.^1^ Conventional methods to study protein-protein interactions involve large-scale affinity purification or separation of protein complexes combined with protein identification using mass spectrometry.^2–7^ These methods typically require ectopic expression of proteins, affinity tags for purification or separation of protein complexes from extracts and other non-physiological conditions, which can produce artifacts and disruption of of native protein interactions. In contrast, cross-linking mass spectrometry (XL-MS) relies on covalent labeling and cross-linking of proteins in physical proximity, thereby identifying physiologic protein-protein interactions. In addition, chemical cross-linking preserves direct or proximal protein-protein interactions, and can also identify transient interactions that occur dynamically during specific cell or developmental states. In addition, high-resolution mass spectrometry can identify direct sites of interaction between cross-linked amino acids, providing detailed spatial information for structural studies.^8–9^ Recently, development of collision-induced dissociation (CID) cleavable cross-linkers^10–14^ and improvements in algorithms for cross-link peptide spectral analysis^15–19^ has enabled protein-protein interaction mapping of complex proteomes, including worm,^20–21^ bacteria,^21–23^ and human^15–16, 24–25^ and various other organisms,^26^ with hundreds-to-thousands of cross-links and protein-protein interactions.^16, 21^ While cross-linking of cell extracts provides direct control of the reaction to label specific protein complexes,^15–16, 20–21^ membrane-permeability of some cross-linking reagents enables cross-linking of protein complexes in live cells to define specific protein-protein interactions *in situ*.

To facilitate large-scale interaction proteomics, we sought to define the optimal methods for the isolation of cross-linked peptides from protein complexes labeled *in situ*, and to ensure their accurate identification using a target-decoy strategy adapted for cross-linking mass spectrometry. Thus, we present optimized parameters for the use of the CID-cleavable cross-linker disuccinimidyl sulfoxide (DSSO) *in situ*. We found that higher-energy collisional dissociation (HCD) fragmentation produced as many cross-linked protein complex identifications as electron transfer dissociation (ETD) fragmentation using either whole proteome or focused target databases for cross-link peptide spectral matching. The use of multiple proteases significantly increased the coverage of protein-protein interactions maps. Importantly, the developed target-decoy strategy allowed for explicit control of false discovery and optimization of sensitivity and specificity of protein interactions maps in cells *in situ*. Application of this approach to human chromatin labeled in live cells recapitulated known and revealed new protein interactions of nucleosomes and other chromatin-associated complexes.

## Experimental Section

### Reagents

Disuccinimidyl sulfoxide (DSSO), formic acid, dimethyl sulfoxide (DMSO), and LC-MS grade solvents were obtained from Thermo Scientific. Bovine serum albumin (BSA), (4-(2-hydroxyethyl)-1-piperazineethanesulfonic acid (HEPES), sodium chloride, magnesium chloride, potassium chloride, sucrose, ethylenediaminetetraacetic acid (EDTA), dithiothreitol, guanidinium hydrochloride, *Staphylococcus aureus* micrococcal nuclease, iodoacetamide, ammonium bicarbonate and formic acid were obtained from Millipore Sigma. Protease inhibitors 4-(2-aminoethyl)-benzenesulfonyl fluoride hydrochloride (AEBSF) and pepstatin were obtatined from Santa Cruz, bestatin from Alfa Aesar and leupeptin from EMD Millipore. Trypsin and GluC proteases were obtained from Promega. Chymotrypsin protease was obtained from Pierce Thermo. LysC was obtained from Wako Pure Chemical Industries.

### Cell culture and protein isolation

HEK293T cells were cultured in DMEM supplemented with 10% (v/v) fetal bovine serum, penicillin and streptomycin. HEK293T cells (50 million) were sedimented by centrifugation at 500 *g* and lysed in 500 μl 6M guanidine hydrochloride in 100 mM ammonium bicarbonate buffer, pH 8. The lysate was sonicated (E210 adaptive focused acoustic sonicator, Covaris) for 5 minutes at 4°C and homogenized by passing through a 27G needle. The lysate was then clarified by centrifugation at 18,000 *g* for 10 min at 4°C. Protein concentration of clarified lysates was determined using the bicinchoninic acid (BCA) assay, according to manufacturer’s instructions (Thermo Scientific).

### Preparation of cross-linked bovine serum albumin and peptide purification

50 μg of BSA was solubilized in 50 μl HEPES buffer (20 mM HEPES, pH 8, 150 mM sodium chloride, 1.5 mM magnesium chloride, 0.5 mM dithiothreitol) and cross-linked by adding 1 μl of 50 mM DSSO dissolved in DMSO (1:1 protein amine:DSSO molar ratio). The solution was incubated at room temperature for 1 hour, before quenching by the addition of 1M ammonium bicarbonate buffer to a final concentration of 20 mM. The cross-linked protein was denatured by adding 50 μl of 6 M guanidine hydrochloride in 100 mM ammonium bicarbonate, pH 8, then reduced by adding 11 μl of 100 mM dithiothreitol at 56°C for 1hr and alkylated with 12 μl of 550 mM iodoacetamide at room temperature for 30 min in the dark. Alkylation was quenched by the addition of 13μl of 100 mM dithiothreitol and incubated at 56°C for 20 min. The solution was diluted with 50 mM ammonium bicarbonate solution, pH 8, for a final concentration of 0.2 M guanidine hydrochloride. For digestion with a single protease, either trypsin, chymotrypsin or GluC was added in 1:50 (enzyme:protein, w/w) concentration and incubated at 37°C for 18 hours. For sequential digestion with LysC followed by trypsin, LysC was first added in 1:100 (enzyme:protein, w/w) concentration and incubated at 37°C for 8 hours. Trypsin was then added in 1:50 (enzyme:protein, w/w) concentration and incubated at 37°C for 18 hours. Digestion was stopped by the addition of formic acid to 1% (v/v) concentration. Peptides were then purified using solid phase extraction with C18 Macrospin columns according to manufacturer’s instructions (Nest Group), and concentrated by vacuum centrifugation.

### Protein cross-linking of live cells in situ and isolation of cross-linked peptides from chromatin

HEK293T cells (50 million) were sedimented by centrifugation at 500 g and then suspended in 1 ml of hypotonic buffer (10 mM HEPES, pH 8, 10 mM sodium chloride, 1 mM magnesium chloride, 0.5 mM dithiothreitol, protease inhibitors). For cross-linking, 50 μl of 50 mM DSSO dissolved in DMSO was added and cells were incubated for 1 hour at 4°C. The reaction was then quenched by the addition of 50 μl of 500 mM Tris-HCl, pH 8. Cells were subsequently subjected to 15 strokes of Dounce homogenization and the solution was then centrifuged for 15 min at 3300 *g* to isolate nuclei. Sedimented nuclei were then resuspended in 500 μl of 0.25 M sucrose, 10 mM magnesium chloride buffer and layered on top of a solution of 500 μl of 0.88 M sucrose, 0.05 mM magnesium chloride. The resultant mixture was centrifuged for 10 min at 1200 *g*. Purified nuclei were then resuspended in 1 ml of 10 mM HEPES, pH 7.4, 0.2 mM magnesium chloride solution and incubated on ice for 10 min to extract nucleoplasm. The solution was then centrifuged for 20 min at 770 *g*. The pellet was next resuspended in 1 ml of 10 mM HEPES, pH 7.4, 500 mM sodium chloride, 0.2 mM magnesium chloride and incubated on ice for 10 min to extract bound proteins. The solution was then centrifuged for 20 min at 770 *g*. Next, the pellet was solubilized in 500 μl of detergent buffer (15 mM HEPES, pH 7.5, 500 mM sodium chloride, 1 mM dithiothreitol, 0.34 M sucrose, 10% glycerol, 0.5% v/v Triton X-100) and incubated on ice for 10 min. The solution was then layered on top of a 500 μl solution of 0.88 M sucrose, 0.05 mM magnesium chloride and centrifuged for 20 min at 3300 *g* to purify chromatin. The chromatin pellet was resuspended in 500 μl of 15 mM HEPES, pH 7.5, 15 mM sodium chloride, 60 mM potassium chloride, 5 mM magnesium chloride, 1 mM calcium chloride, 0.25 M sucrose and protease inhibitors (0.5 M AEBSF, 0.001 mM pepstatin, 0.01 mM bestatin and 0.1 mM leupeptin). The chromatin suspension was incubated at 37°C for 10 min. Five units of micrococcal nuclease were subsequently added and incubated at 37°C for 1 hour. The nuclease digestion was stopped by the addition of EDTA to a final concentration of 50 mM and the solution was centrifuged for 10 min at 18,000 *g* to remove the nuclear matrix. The supernatant containing cross-linked chromatin was then quantified using the BCA assay.

Cross-linked chromatin proteins were then purified by filter-aided sample preparation (FASP) using Sartorius Vivacon 500 with a 30,000 MW cut-off filter, as previously described.^27–28^ Briefly, 200 μg of protein was diluted 5-fold in urea buffer (8 M urea, 50 mM ammonium bicarbonate, pH 8.0) and applied to the filter unit. Filter units were centrifuged at 14,000 *g* for 20 min. The membrane was then washed twice with 200 μl urea buffer by centrifugation at 14,000 *g* for 20 min. The membrane was then incubated with 200 μl of 100 mM dithiothreitol in urea buffer for 20 min at room temperature before centrifugation at 14,000 *g* for 20min. The proteins were then incubated with 200 μl of 100 mM iodoacetamide in urea buffer for 20 min at room temperature in the dark before centrifugation at 14,000 *g* for 20 min. The membrane was washed twice more with urea buffer then once with 200 μl of 50 mM ammonium bicarbonate, pH 8. Proteins retained on the membrane were then digested with trypsin at 1:50 (enzyme:protein, w/w) concentration at 37°C for 18 hours. Digested peptides were collected by centrifugation at 14,000 *g* for 20 min. Peptides were then purified using solid phase extraction with C18 Macrospin columns according to manufacturer’s instructions (Nest Group). The purified peptides were concentrated by vacuum centrifugation and resuspended in 100 μl of 5% acetonitrile, 0.1% formic acid. The peptides were then injected onto the strong cation exchange (SCX) column (Protein Pak Hi-Res SP 7μm 4.6 × 100mm, Water) at a flow rate of 0.5 ml/min with a column temperature of 30°C using the Alliance HPLC system (Waters). Gradient was run with 5% acetonitrile and 0.1% formic acid in water for solvent A and 1 M potassium chloride in 5% acetonitrile and 0.1% formic acid in water for solvent B. Column gradient was set as follows: 0-3 min (0% B); 3-33 min (0-10% B); 33-43 min (10-100% B); 43-60 min (100% B); 60-70 min (100-0% B); 70-90 min (0% B). Six fractions were collected from 38 to 60 min and purified using solid phase extraction with C18 Macrospin columns according to manufacturer’s instructions (Nest Group). Peptides were concentrated by vacuum centrifugation and then stored at −20°C before analysis.

### Liquid chromatography and nanoelectrospray mass spectrometry

Cross-linked peptides were separated by reverse phase nanoflow liquid chromatography (EKspert nanoLC 425, Ekisgent) coupled to the Orbitrap Fusion mass spectrometer (Thermo). Cross-linked BSA peptides and HEK293T peptides were resuspended in 0.1% formic acid in water (v/v). For cross-linked BSA peptides, 1 μg of peptide was injected for analysis. For cross-linked BSA peptides mixed with non-cross-linked HEK293T peptides in 1:1 (w/w) ratio, 1 μg of total peptide was analyzed. For SCX fractions of cross-linked chromatin, peptides were resuspended in 20 μl of 0.1% formic acid in water (v/v) and 0.5 μg of peptides were analyzed.

For liquid chromatography, a trap-elute system was used, as described previously.^29^ Reverse phase columns were fabricated as previously described.^30^ Trap columns were fabricated by packing Poros R2 10μm C18 particles (Life Technologies) into 4 cm fritted capillaries with internal diameters of 150 μm. Reverse phase columns were fabricated by packing Reprosil 1.9 μm silica C18 particles into 40 cm fritted capillaries with internal diameters of 75 μm. Peptides were resolved over a 90 min gradient from 5% to 40% acetonitrile and ionized using the DPV-565 Picoview ion source (New Objective) operated with the ionization voltage of 1700 V. Custom electrospray emitters were fabricated as previously described.^31^

For mass spectrometry, four different acquisition methods were used: 1) CID-MS2/HCD-MS2, 2) CID-MS2/ETD-MS2, 3) CID-MS2/EThcD-MS2, 4) CID-MS2/HCD-MS3, where HCD is the higher-energy collisional dissociation, ETD is electron transfer dissociation, and EThcD is the hybrid electron transfer with supplemental activation by higher energy collision dissociation. For MS2-MS2 methods, precursor ion spectra were recorded at 400-1800 *m*/*z* with 60,000 *m*/*z* Orbitrap resolution, automatic gain control target of 1 ×10^5^ ions and maximum injection time of 50 ms. Precursor ions with 3-10 positive charge were selected for MS2 fragmentation with dynamic exclusion of 60 secs after 1 scan and isolation window of 2 Th. For CID-MS2, precursor ion spectra were recorded in the Orbitrap with resolution 30,000 m/z, automatic gain control target of 5.0 ×10^4^ and maximum injection time of 100 ms at normalized collision energy of 30%. For HCD-MS2, precursor ion spectra were recorded in the Orbitrap with resolution 30,000 m/z, automatic gain control target of 5.0 ×10^4^ and maximum injection time of 120 ms at normalized collision energy of 30%. For ETD-MS2, precursor ion spectra were recorded in the Orbitrap with resolution 30,000 m/z, automatic gain control target of 5.0 ×10^4^ ions and maximum injection time of 120 ms with fluoroanthene reaction time of 50 ms, ETD target of 2.0 ×10^5^ ions and maximum fluoroanthene injection time of 200 ms. For EThcD-MS2, precursor ion spectra were recorded in the Orbitrap with resolution 30,000 m/z, automatic gain control target of 5.0 ×10^4^ ions and maximum injection time of 120 ms with reaction time of 50 ms, reagent target of 2.0 ×10^4^ and maximum fluoroanthene injection time of 200 ms and supplemental collision energy of 20%. For HCD-MS3, MS2 fragment ions with 2-6 charge states and mass difference of 31.9721 Da with 1-100% intensity range were selected for MS3 fragmentation, with a mass tolerance of 30 ppm and 2 Th isolation window. MS3 fragment ion spectra were recorded in the linear ion trap acquisition with automatic gain control target of 2.0 ×10^4^ ions and maximum injection time of 150 ms.

### Data analysis

Acquired mass spectra raw files were analyzed using XlinkX version 2.2,^15–16^ as implemented in Proteome Discoverer version 2.2. The target search database was the *Homo sapiens* proteome from SwissProt, containing isoforms^32^ (version January 2016), supplemented with the sequences of common contaminant proteins from cRAP.^33^ Mass spectral analysis parameters included 2 maximum allowed missed cleavages, minimum peptide length of 5 and 4 maximum variable modifications: methionine oxidation, hydrolysis of lysyl-DSSO by water, and lysyl-DSSO-Tris adduct. Precursor mass tolerance was set at 10 ppm with fragment mass tolerance of 20 ppm and 0.5 Da for Orbitrap and ion trap spectra, respectively. Cysteine carbamidomethylation was set as fixed modification. A 1% false discovery rate (FDR) was set for cross-link peptide spectra matching with minimum score threshold set at 0 using the Percolator algorithm.^34^ Peptide spectral matches and their scores were analyzed using R scripts to concatenate cross-links identified from biological replicates (https://github.com/kentsisresearchgroup/R-script-for-XLMS-data-analysis). For histone proteins, peptides that could be identified as exclusively belonging to specific isoforms were marked with their specific isoform protein name or gene name. Proteins that could be mapped to multiple isoforms were labeled with the family protein or gene name, e.g. peptides that could belong to histone isoforms H3.1, H3.2 or H3.3 were labeled as histone H3 peptides. Bar graphs and scatter plots were plotted using Origin 2018 (Microcal). Venn diagrams were made with Venny version 2.1.^35^ Peptide cross-link sequences were visualized using xiNET.^36^ Atomic resolution structures were visualized with UCSF Chimera^37^ version 1.12 and cross-links were mapped using Xlink analyzer.^38^ Protein-protein interaction maps were made with Cytoscape version 3.6.0.^39^ Mass spectrometry raw files and search results are publicly available through the ProteomeXchange data repository via the PRIDE database.

## Results and Discussion

### Calibration of cross-link peptide-spectra matching (CSM) scores to control sensitivity and specificity of large-scale XL-MS

MS-cleavable cross-linkers are designed to improve the accuracy and sensitivity of identification of protein-protein interactions by mass spectrometric analysis and subsequent statistical spectral matching. Specifically, the DSSO cross-linker is labile upon collision induced dissociation, leading to decoupling of cross-linked peptides into linear peptides in the gas-phase, and consequently enabling independent identification of each peptide. In addition, fragmentation of the sulfoxide cross-linker leaves characteristic adducts on the de-coupled peptides, thus producing diagnostic mass differences that can be leveraged to improve the specificity of identification of cross-linked peptide ions^12^. Depending on the experimental parameters used, fragmentation of the peptide backbone can then be achieved by selecting either the cross-linked precursor ion or the decoupled peptide ions. Algorithms based on scoring functions for matching of cross-linked peptide spectra have recently been described.^15–16, 19^

To further advance our ability to control the sensitivity and specificity of cross-linked peptide discovery, we reasoned that discrimination of true and false positive peptide-spectral matches can be achieved using target-decoy strategies tailored specifically to cross-linked peptides. To model a typical XL-MS sample, containing both cross-linked and non-cross-linked peptides, we cross-linked purified bovine serum albumin (BSA), which was subsequently diluted in non-cross-linked proteins extracted from human HEK293T cells (Figure 1A). BSA is a widely used model protein for mass spectrometry that spontaneously oligomerizes in solution.^40–41^ Insofar as HEK293T cells do not express human albumin or proteins with sequence homology to BSA, this mixture model of true positive (cross-linked BSA) and true negative (human proteins) should provide a faithful model to estimate and control the FDR of large-scale cross-linked mass spectrometry. Thus, we use a generalized linear mixture model to estimate the false discovery rate among four different methods of cross-linked peptide analysis: CID-MS2/HCD-MS2, CID-MS2/ETD-MS2, CID-MS2/EThcD-MS2 and CID-MS2/HCD-MS3 (Figure 1B). Despite applying a 1% FDR threshold for peptide-spectral matching using Percolator,^34^ we observed a substantial number of apparently incorrectly matched cross-linked peptide spectra. For example, for CID-MS2/HCD-MS2 fragmentation, 173 of 291 (59%) of cross-links matched were mapped to human proteins that were not DSSO cross-linked, indicating incorrect CSMs. Manual inspection of representative spectra confirmed that these apparent incorrect matches were due to either incomplete fragmentation spectrum of one of the peptides in the pair or relatively low signal intensity of the diagnostic DSSO fragment ions. We confirmed that none of the examined spectra from apparently incorrect matches exhibited fragment ions consistent with albumin peptide sequences. Finally, we also verified that none of the mis-matched spectra were due to the incomplete DSSO cross-linking reaction due to spectra with single site DSSO adducts.

**Figure 1.**
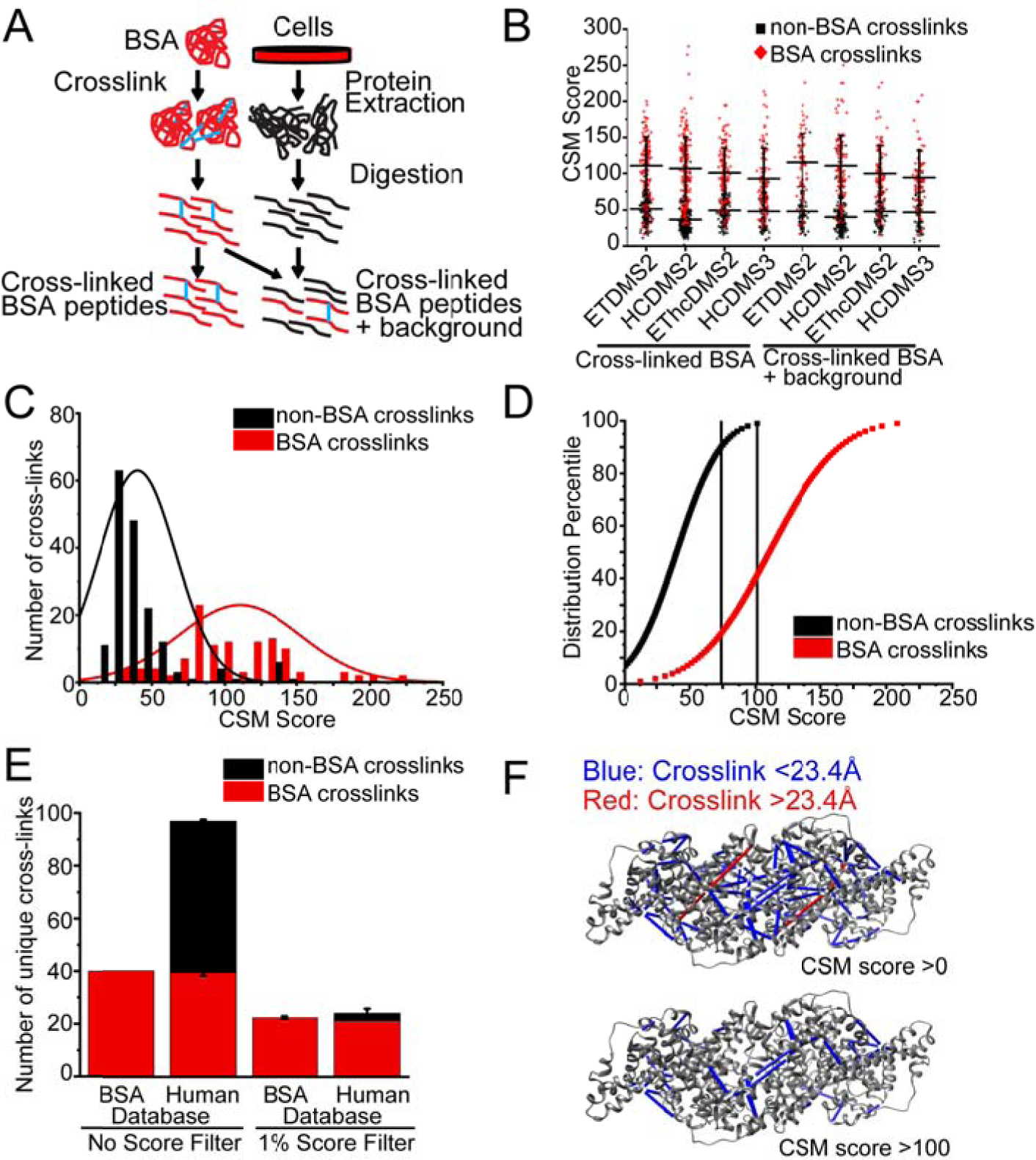
Calibration of cross-link peptide-spectra matchi g (CSM) scores to control sensitivity and speci icity of large-scale XL-M. A) Workflow for preparation of cross-linked BSA peptides and cross-linke BSA peptides spiked into non-cross-linked proteome background peptides. B) High number of BSA cross-linked peptides (red) and on-BSA cross-linked peptides (black) identified across fragmentation meth ds and in both cross-linked BSA and cross-linked BSA spiked into background. C) Fitted Gaussian distribution of cross-link spectra matching score of non-BSA cross-links and BSA cross links for cross-linke BSA only sample acquired with CID-MS2/HCD-MS2 fragmentation. D) Percentile plot of gaussi n distribution of CS score with 1% and 10% score filter to liminate 99% and 90% of non-BSA cross-links. E) Comparison of number of cross-links identified when no score filter is applied and when 1% score filter is applied F) Cross-links violating physical distance constraint of cross-linker (red) are eliminated when 1 score filter is applied when mapping cross-links to crystal structure (PDB ID 4f5s).

We reasoned that such false cross-linked peptide-spectral matches can be distinguished by their relative cross-link peptide-spectra matching (CSM) scores, as defined by the current XlinkX algorithm (Figure 1B). ^15–16^ Thus, we fit the observed CSM score distributions to a linear mixture of two Gaussian functions, producing two modes corresponding to apparently true and false matches (Figure 1C). Consequently, for the observed spectra using the four acquisition methods tested, this led to a CSM score threshold of 100 to achieve 1% FDR at the peptide level (Figure 1D). In addition, we defined a CSM score threshold of 74 for improved sensitivity at a more relaxed (10%) peptide-level FDR. These empiric CSM score thresholds should be generalizable to other experiments, insofar as the physicochemical properties of the peptide analyzed are adequately approximated by BSA and human proteins (Figures S-1).

Importantly, the use of these CSM score thresholds reduced the number of false non-BSA cross-links matched to non-cross-linked human proteins from 58 to 3 (Figure 1E). In addition, we analyzed the identified BSA cross-links with respect to the high-resolution atomic-level structure of BSA (PDB ID 4f5s).^42^ Consistently, the use of CSM score thresholds eliminated cross-links between lysine residues that are farther than 23 Å (the length of the DSSO cross-linker), while retaining cross-link matches with the expected physical distances (Figure 1F). These results demonstrate the need for controlling the false identifications inherent in the statistical spectral matching for large-scale cross-linking mass spectrometry proteomics, and provide a facile means to do so using a target-decoy spectral matching strategy adapted for cross-linking mass spectrometry.

### Improved cross-linked peptide identification using CID-MS2/HCD-MS2 fragmentation

Comparison of CID, HCD and ETD fragmentation of linear non-cross-linked peptides indicates that the efficiency of each fragmentation method, and therefore their capacity to produce optimal fragment ion spectra, depends on the specific physicochemical properties of fragmented peptides^43^. For example, ETD exhibits reduced fragmentation efficiency for larger *m/z* peptides.^44^ Previous studies of non-cleavable cross-linkers found that HCD fragmentation increased the identification of cross-linked peptides with charge states of 3-4, while the hybrid EThcD method facilitated the identification of cross-linked peptides with charge states of ≥5.^45^ Thus, to assess the various methods of identification of DSSO cross-linked peptides, we used the mixture of cross-linked BSA and non-cross-linked human proteome. We compared 4 different fragmentation methods, with CID-MS2 used for cross-link fragmentation: CID-MS2/HCD-MS2, CID-MS2/ETD-MS2, CID-MS2/EThcD-MS2 and CID-MS2/HCD-MS3.

We observed an average of 43 cross-links per experiment across 3 biological replicates identified by CID-MS2/HCD-MS2, as compared to 25, 19 and 7 cross-links identified with CID-MS2/ETD-MS2, CID-MS2/HEThcD-MS2 and CID-MS2/HCDMS3 methods, respectively (Figure 2A). A total of 56 unique cross-links across 3 biological replicates were identified using CID-MS2/HCD-MS2 fragmentation, while 35 unique cross-links were identified from CID-MS2/ETD-MS2, of which only 3 unique cross-links identified by CID-MS2/ETD-MS2 were not identified by CID-MS2/HCD-MS2 (Figure 2B). For cross-linked BSA peptides mixed with the non-cross-linked proteome, an average of 21 cross-links per experiment across 3 biological replicates were identified by CID-MS2/HCD-MS2, as compared to 19, 13 and 3 cross-links identified by CID-MS2/ETD-MS2, CID-MS2/HEThcD-MS2 and CID-MS2/HCDMS3 methods, respectively (Figure 2C). 28 unique BSA cross-links across 3 biological replicates were identified from CID-MS2/HCD-MS2, while a comparable 27 unique cross-links were identified from CID-MS2/ETD-MS2. Both CID-MS2/HCD-MS2 and CID-MS2/ETD-MS2 identified 8 and 5 cross-linked peptides, not identified by other fragmentation methods, respectively, suggesting that there is some orthogonality in coverage between these two fragmentation methods (Figure 2D). Comparison between CID-MS2/HCD-MS2 and CID-MS2/ETD-MS2 methods showed an overlap between the types of cross-links identified, with 32 out of 59 (54%) and 19 out of 35 (54%) shared cross-links identified for cross-linked BSA peptides and cross-linked BSA peptides mixed with the non-cross-linked proteome, respectively. Comparison of identified cross-linked peptides showed relatively shorter peptide length, lower precursor *m/z* and lower cross-linked peptide identifications across all charge states obtained by ETD fragmentation compared to that obtained by HCD fragmentation (Figure S-1), in line with prior observations.^44^ A substantial number (10-30%) of cross-linked peptides were identified exclusively by either ETD or HCD fragmentation methods (Figure 2B, 2D), suggesting some orthogonality between the two fragmentation methods. Naturally, these results are generalizable to other complex proteomes largely for cross-linked peptides that are adequately modeled by cross-linked BSA.

**Figure 2.**
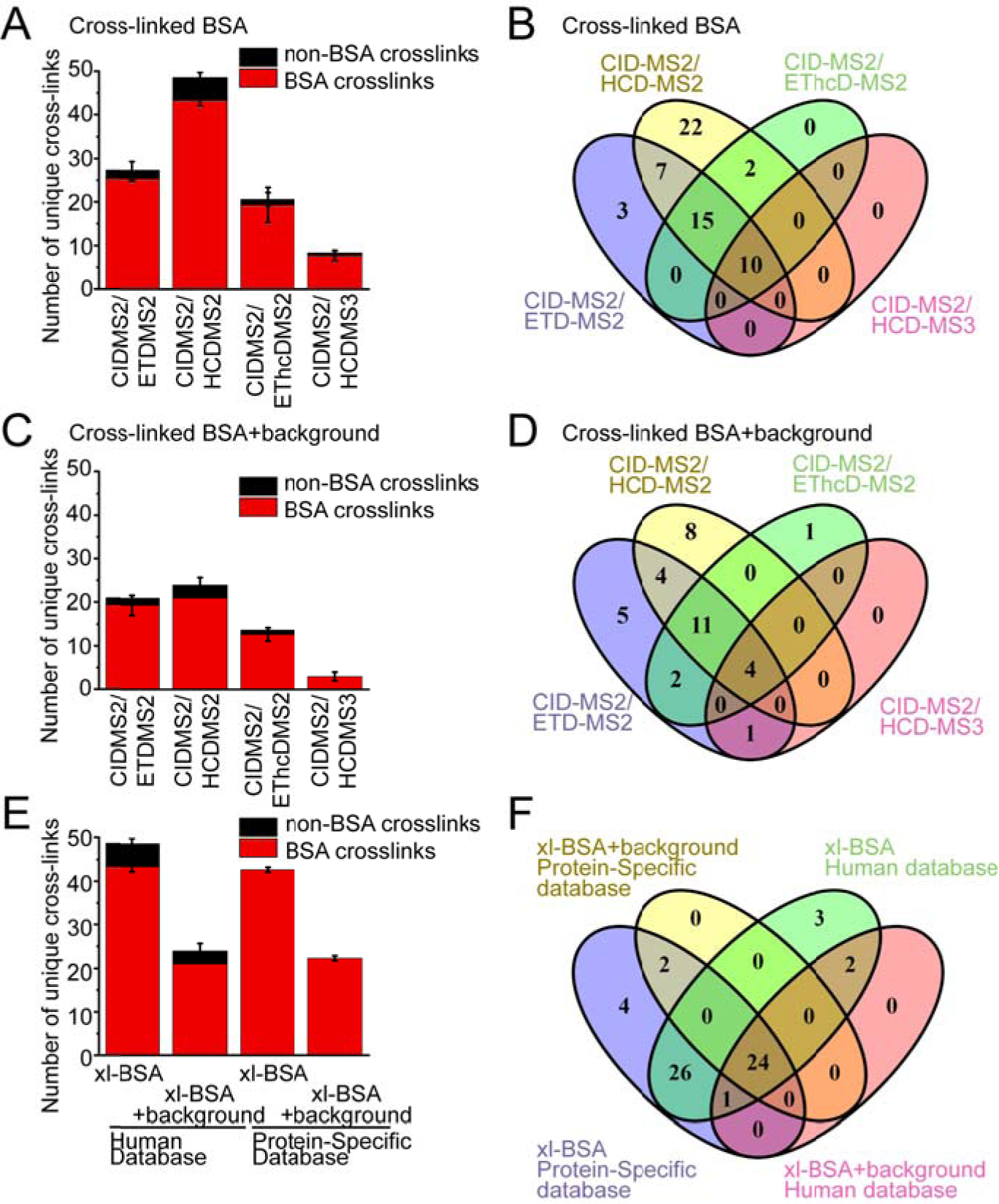
I proved cross-linked pepti e identification using CID-MS2/HCD-S2 fragmentation. Comparison of 4 different fr gmentation methods: CID-MS2/HCD-S2, CID-MS2/ETD-MS2, CID-MS2/E hcD-MS2 a d CID-MS2/HCD-MS3 and data ases: BSA protein specific database or human proteome database with BSA for increasing cross-link p ptide identifiation. A) Comparison between 4 fragmentation methods for averag number of cross-link identificatio s of cross-linked BSA sam le. BSA cross-links colored red and non-BSA cross-links colored black. Error bars represent standard de iation from three biological replicates. B) Comparison etween 4 fragmentation methods for overlap of all unique cross-link identific tions C) Comparison between 4 fragmentation methods for average number of cross-link i entifications of cross-linked BSA pepti es spiked into human proteome background. D) Comparison between 4 fragmentation ethods for overlap of all unique cross-link identifications of cross-linked BSA sample spiked into human proteome background. E) Comparison bet een 2 databases for average number of cross-link identifications of cross-linked BSA peptides and cross-linked BSA p ptides spiked into human proteome back round. F) Comparison between 2 databases for overlap of all unique cross-link identifications of cross-linked BSA peptides and cross-linked BSA peptides spiked into human proteo e background.

In all, these results indicate that CID-MS2/HCD-MS2 fragmentation can yield the highest number of cross-links for simple samples, with a 60% increase in unique cross-link sites as compared to CID-MS2/ETD-MS2. For complex samples, CID-MS2/HCD-MS2 identified as many unique cross-link sites as CID-MS2/ETD-MS2. On the other hand, CID-MS2/HCD-MS3 and CID-MS2/EThcD-MS2 methods consistently recorded fewer cross-link spectra and fewer identifications when compared to CID-MS2/ETD-MS2 and CID-MS2/HCD-MS2 methods. This can be attributed to the relatively longer acquisition times required for MS3 and EThcD methods, as also noted in a recent study.^16^

### Sample-specific search target databases for identification of protein-protein interactions by XL-MS

While the use of a mass spectrometer cleavable cross-linkers reduces the database search space and time as compared to non-cleavable cross-linkers,^15^ the use of sample-specific search databases can improve the sensitivity of mass spectral matching.^46–49^ Since cross-linking mass spectrometry is often used for the analysis of specific purified protein complexes, we examined the effects of using sample-specific as compared to proteome-wide search databases.

Thus, we generated search databases based on the BSA sequence alone, or a concatenated database containing BSA plus the human reference proteome, supplemented with common proteomic contaminants from cRAP.^33^ We observed an average of 43 BSA cross-links across 3 biological replicates when searching against either the BSA-specific or reference human proteome databases (Figure 2E). For cross-linked BSA peptides mixed with the non-cross-linked proteome, we found an average of 21 and 22 BSA cross-links from 3 biological replicates when searching against the BSA-specific database or BSA plus reference human proteome databases, respectively (Figure 2E). This suggests that the database size and composition do not affect the sensitivity of cross-link identification *per se* (Figure 2F). With the BSA-specific database excluding any possible false matches, we observed a significant reduction in false matches of cross-linked BSA to non-cross-linked human proteins, when using the BSA-specific database, as would be expected from first principle considerations (0 as compared to 3, respectively, *p* = 0.048, Figure 2E).

### Diverse sequence-specific proteases to expand the coverage and density of DSSO XL-MS protein interaction maps

We reasoned that the DSSO reaction with lysine residues reduces the accessibility of cross-linked peptides for protease digestion, thus reducing the identification of tryptic cross-linked peptides. Consistent with this, we found substantially more unique cross-links identified from the sequential protease digest using LysC endoprotease followed by trypsin, as compared to that of trypsin alone (90 versus 18, respectively, Figure 3A). This combined LysC and trypsin digestion produced cross-link sites across the whole BSA protein sequence (Figure 3C), suggesting that accessibility of amino acid sites for protease digestion affects cross-link peptide identification. For linear non cross-linked peptides, additional sequence-specific proteases are known to identify additional peptide sequences, thus improving protein sequence coverage.^50–52^ Hence, we compared three different digestion methods using additional proteases specific to non-lysine amino acids: 1) sequential digestion with LysC followed by trypsin, 2) GluC endoprotease, and 3) chymotrypsin. Combined LysC and trypsin digestion identified the highest number of unique cross-linked sites, as compared to GluC and chymotrypsin digestions (90 versus 69 and 56, respectively, *p* < 0.01 and *p* < 0.01, Figure 3B). We observed a minor overlap among these cross-linked sites, with the majority of identified sites involving different BSA residues (Figure 3C), presumably due to the differences in susceptibility to enzymatic endoproteolysis.^50^ On average, cross-linked peptides identified by GluC digestion were more highly charged, as compared to those identified by chymotrypsin or trypsin (Figure S-2). Thus, the use of multiple, non-lysine-dependent proteases can expand the coverage and density of DSSO XL-MS protein interaction maps.

**Figure 3.**
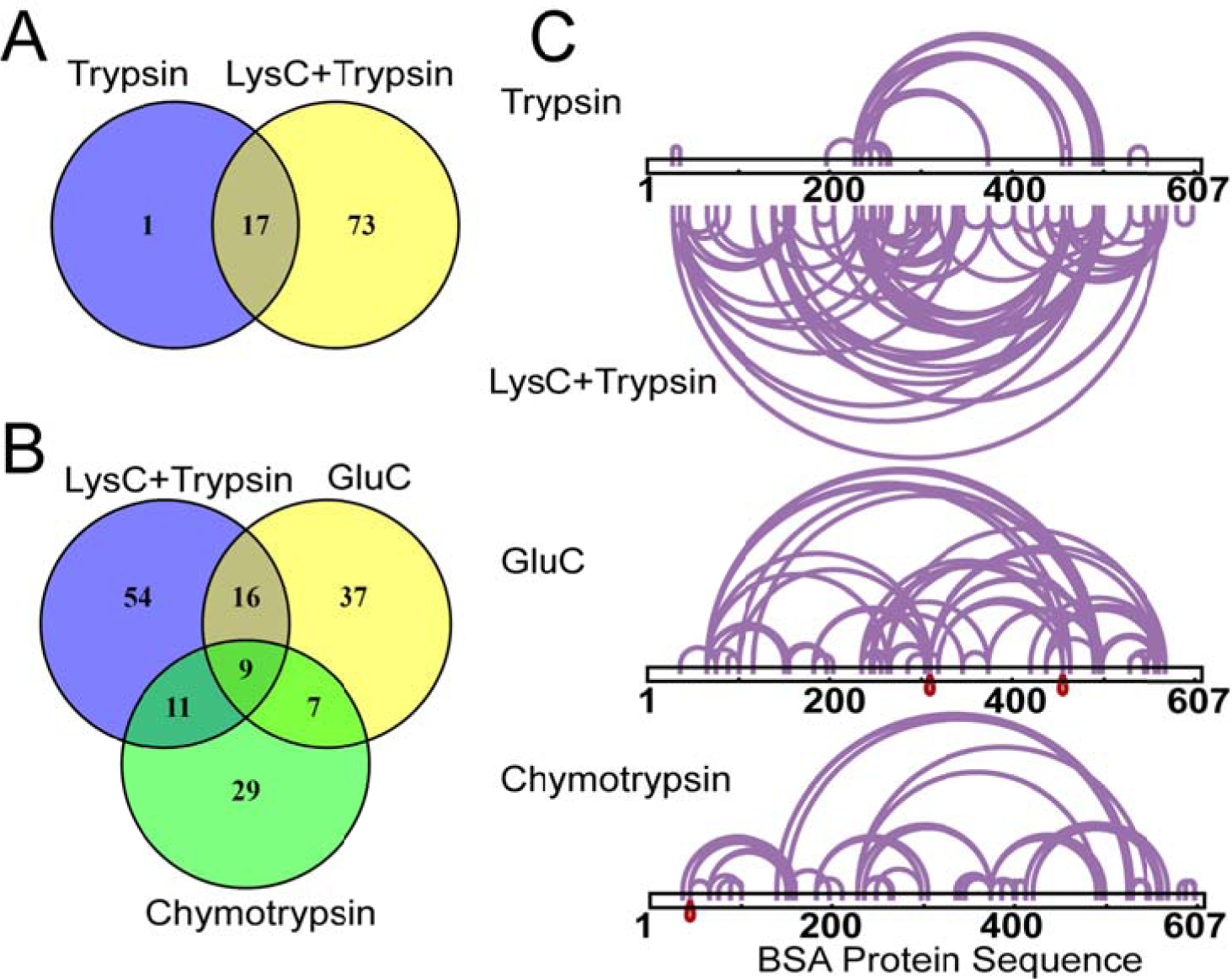
Mul iple proteases to expand the coverage nd density of DSSO XL-MS protein interaction maps. A) Combined LysC and trypsin protease digestion greatly increases number of cross-link pept des identified compared to trypsin only protease digestion, B) Low number of overlapping cross-linked peptides identified from LysC and trypsin or GluC only or chymotrypsin only protease igestion, C) Visualization f BSA cross-link sites identified from trypsin only, LysC and trypsin, GluC only or chymotry sin only protease digestion, as analyzed using CID-S2/HCD-MS2 fragmentation.

### Native chromatin protein-protein interactions identified using cross-linking mass spectrometry *in situ*

Having identified a robust means to control the sensitivity and specificity of large-scale cross-linking mass spectrometry, we sought to examine the ability of DSSO to label native protein complexes in live human cells. We chose to study protein-protein interactions of human chromatin *in situ*, given its abundance and prominent functions in the regulation of cell growth and development. We optimized a protocol to label live human HEK293T cells with DSSO (see Methods). Chromatin proteins were extracted from purified nuclei by sucrose sedimentation, followed by sequential salt, detergent, and micrococcal nuclease digestion to release chromatin bound proteins.^53–55^ We used SDS-PAGE followed by silver staining to confirm the presence of reduced mobility protein complexes upon DSSO cross-linking, as compared to mock-treated cells (Figure 4A). Likewise, we used Western immunoblotting to verify the isolation of histone-containing chromatin complexes, as opposed to nucleoplasmic or cytoplasmic proteins, as assessed by histone H3, BRG1, and GAPDH, respectively (Figure 4B).

**Figure 4.**
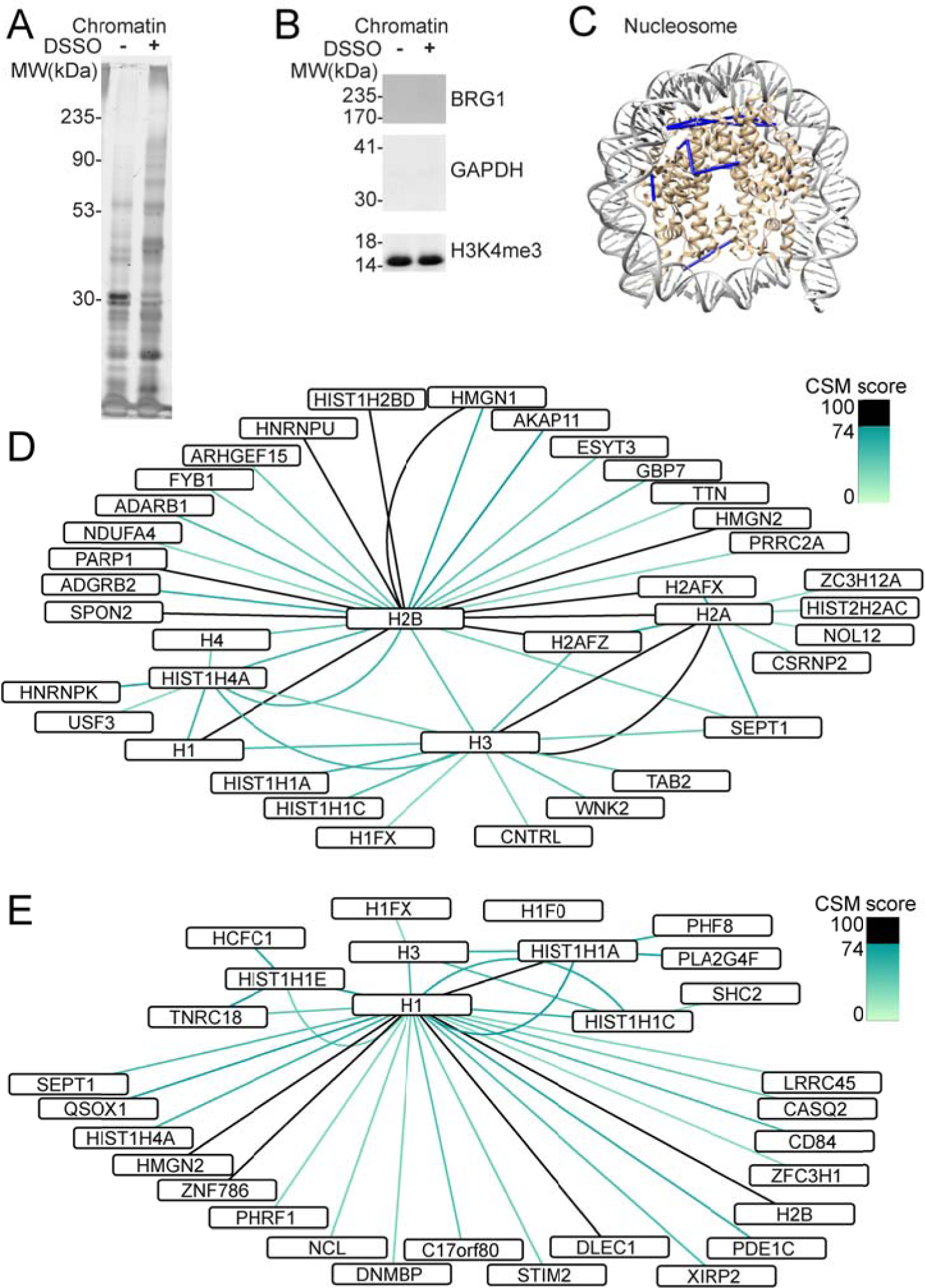
Native chr matin protein-protein interactions identified using cross-linking mass spectrometry *in situ*. A) Purified chromatin bound proteins are cross-linked by cross-linking *in situ*. B) Enrichment of histone protein (H3K me3) with absence of nuclear protein (B G1) and cytoplasmic protein (GAPDH) from chromatin bound proteins by western blot. C) Cross-links **i**dentified map onto known high resolution nucleosome structure (PDB ID 3av1). D) Map of protein-protein interactions between core histone proteins (H2A, 2B, H3, H4) colored accor ing to CSM score. E) Map of protein-protein interactions involving linker histone H1.

Analysis of the cross-linked chromatin fraction identified 1,277 unique cross-linked peptides from three biological replicates, with 384 unique cross-linked peptides at least two out of three biological replicates (Table S-1). Of these, 189 (49%) of cross-linked peptides were between two different proteins, while 195 (515) were interactions within the same proteins, with 95 of these interactions also represented in the current BioGRID and CORUM protein-protein interaction databases.^56–59^ In agreement with the empirically determined CSM score thresholds for identifying high-confidence protein-protein interactions, we observed that all 8 cross-links involving residues in the nucleosome histone core can be mapped onto the high-resolution, atomic-level structure of the human nucleosome,^60^ consistent with the 23 Å distance constraint based on the length of the DSSO linker (Figure 4C). Likewise, protein-protein interactions involving the core histone proteins captured many known histone-histone interactions, such as those between the core histone H2B and H2A, H4 and H3 histones (Figure 4C). Notably, we observed 46 cross-links involving the histone tails, which were not visualized in the isolated nucleosome core structure *in vitro*,^60^ suggesting that they are in fact organized and bound to specific cofactors *in situ*. In particular, we observed numerous cross-links involving histone H2B K5 and histone H3 K4 and K18, which were also found in a recent study.^61^

Notably, using labeling of protein complexes *in situ*, we observed numerous previously unrecognized interactions. For example, we found numerous cross-links between the HMGN1 and HMGN2 proteins with histone H2B and histone H1.2 (Figure 5D & E). HMGN1 and HMGN2 are known to regulate chromatin configuration,^62^ which can involve phosphorylation of histone H2A and histone H3.^63–64^ Our findings indicate that HMGN1 can also bind directly to K120 of H2B, and HMGN2 can bind directly to K206 of H1.2 in the context of native chromatin *in situ*, which may provide the sought-after mechanism to explain its structural effects on chromatin configuration directly.^65–66^ Likewise, we observed new protein-protein interactions involving ADARB1 with histone H2B, and centriolin/CNTRL with histone H3. This suggests that these two proteins may have unanticipated functions on DNA and/or chromatin. Similarly, we observed numerous cross-links to the linker histone H1, including PHRF1, ZNF786, PDE1C, and SEPT1, suggesting that these proteins may have additional structural chromatin functions, possibly in stabilizing or regulating histone linker-dependent higher-order chromatin conformations *in situ* (Figure 4E).

**Figure 5.**
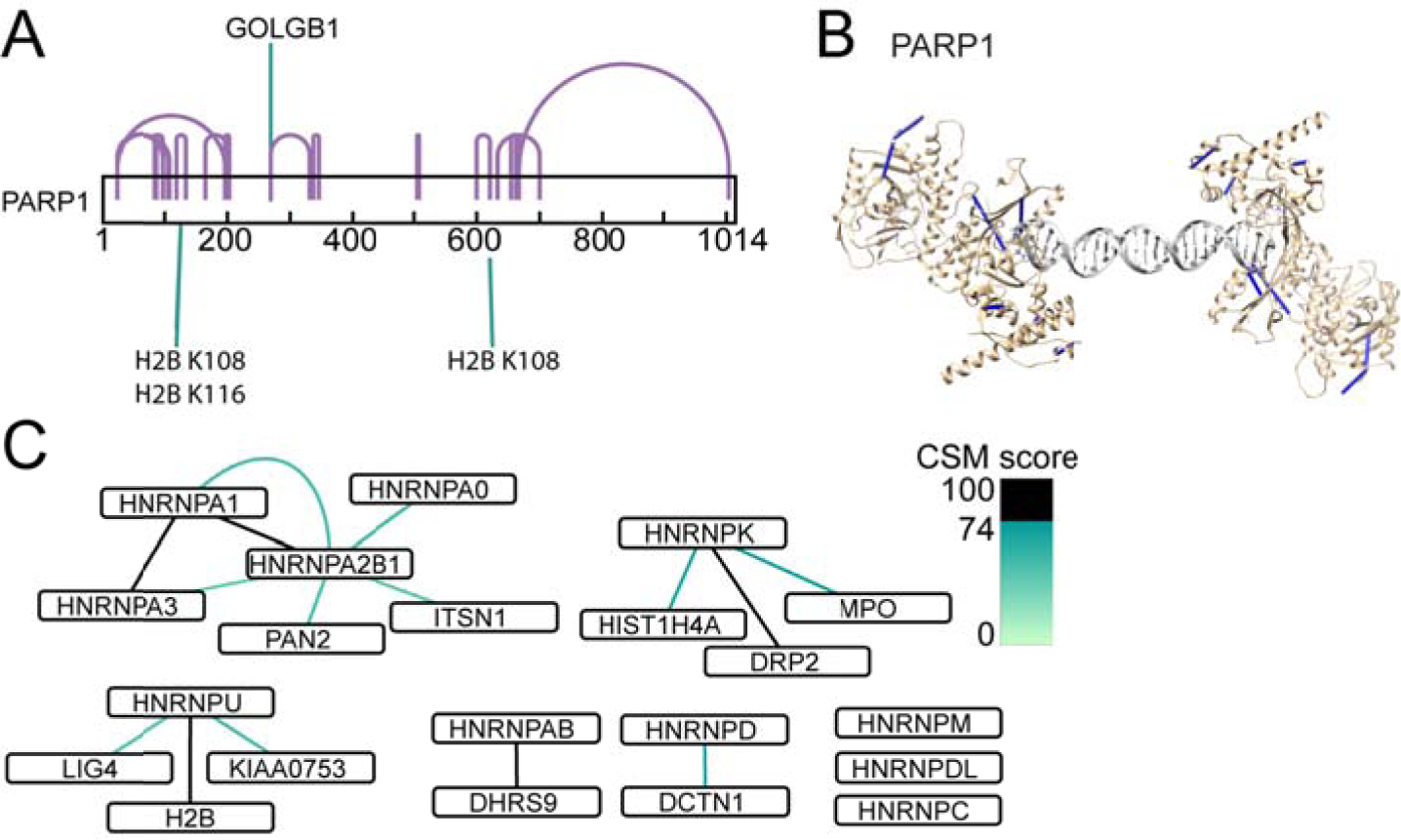
Chromatin interactions of proteins involved in DNA damage repair and ribonucleoproteins. A) DNA repair protein PARP1 interacts with Histone H2B. B) PARP1 cross-links map nto known high resolution structure of PARP1 bound to DNA double stranded break (PDB ID 4dqy). C) Protein-protein interactions of RNA-binding heterogene us nuclear ribonucleoprotei s (hnRNPs).

Lastly, we found numerous chromatin interactions of proteins involved in DNA damage repair and ribonucleoproteins. For instance, we observed cross-links from PARP1 to K108 and K116 of histone H2B, in addition to 17 intra-protein PARP1 cross-links (Figure 5A). We confirmed that 8 of 17 detected PARP1 cross-links could be mapped onto its high-resolution structure (PDB ID 4dqy),^67^ as bound to a DNA double stranded break site (Figure 5B). PARP1 is known to self-associate,^68–69^ and our findings provide a direct mechanism of chromatin recruitment, linking PARP1 to histone H2B. This interaction may affect PARP1 self-association and DNA recognition and/or contribute to chromatin remodeling associated with DNA damage repair.^70–74^ Similarly, we found numerous cross-links among the heterogenous nuclear ribonucleoproteins hnRNPs and chromatin (Figure 5C). This included known interactions between hnRNPA1 and hnRNP2B, and between hnRNPA1 and hnRNPA3, providing specific structural details on their physical proximity.^75–76^ In addition, we observed direct interactions between DNA ligase LIG4 and hnRNPU proteins with histone H2B, raising the possibility that chromatin and hnRNP recruitment can regulate repair of DNA double-strand breaks.^77^ While these interactions will require dedicated future studies to confirm and establish their functions, these results indicate that native chromatin protein-protein interactions can be identified by using cross-linking mass spectrometry *in situ*.

## Conclusions and Future Directions

Here, we sought to define the optimal methods for the isolation of cross-linked peptides from protein complexes labeled *in situ*, and to ensure their accurate identification using a target-decoy strategy adapted for cross-linking mass spectrometry. The presented generalized linear mixture model offers a facile means to control sensitivity and specificity based on a target-decoy spectral matching strategy adapted for cross-linking mass spectrometry. Using this approach, additional mixtures can be prepared in the future to model distinct protein complexes and peptide sequences, such as those containing particular chemical modifications. We also anticipate that improved scoring functions incorporating fragmentation features of specific cross-linkers as a function of fragmentation and isolation methods, such as the diagnostic sulfoxide fragment ion doublet in DSSO, as well as the use of isotopically labeled heavy and light cross-linkers to label experimental and control samples, can be used to improve the sensitivity and specificity of cross-linked peptide spectral identification.^78–80^ Likewise, for the study of specific complexes, sample-specific target databases can be used to improve spectral matching accuracy. Importantly, we found that the density and coverage of protein-protein interaction maps can be significantly improved through the use of diverse sequence-specific proteases, and complex cross-linked mixtures can be effectively analyzed using relatively fast CID-MS2/HCD-MS2 fragmentation methods. Lastly, our studies of native human chromatin labeled in live cells recapitulated known and revealed new protein interactions of nucleosomes and other chromatin-associated complexes. In all, this approach should facilitate the discovery and definition of protein-protein complexes *in vivo*, both in health and disease.

## Acknowledgements

We thank Rosa Viner and David Horn for support of the beta version of XLinkX and Yael David for comments on the manuscript. This work was supported by the NIH R01 CA204396, P30 CA008748, Burroughs Wellcome Fund, Josie Robertson Investigator Program, Rita Allen Foundation, Alex’s Lemonade Stand Foundation, St. Baldrick’s Arceci Innovation Award, (A.K.), and Agency for Science, Technology and Research, Singapore (Z.S.). A.K. is the Damon Runyon-Richard Lumsden Foundation Clinical Investigator.

## Notes

The authors declare no competing financial interest.

## Supporting Information

**Figure S-1.** Characterization of peptide length, m/z and charge state of cross-linked peptides from comparison of fragmentation methods: CID-MS2/HCD-MS2, CID-MS2/ETD-MS2, CID-MS2/EThcD-MS2 and CID-MS2/HCD-MS3. A) Cross-link peptide length by number of amino acids from cross-linked BSA peptide spectra. B) Precursor *m/z* values. C) Number of cross-linked peptide spectra based on charge state. D,E,F,G) Comparison of peptide A and peptide B length from cross-linked peptides identified by CID-MS2/HCD-MS2, CID-MS2/ETD-MS2, CID-MS2/EThcD-MS2 or CID-MS2/HCD-MS3.

**Figure S-2.** Characterization of peptide length, m/z and charge state of cross-linked peptides from comparison of multiple protease digestion: Trypsin only, combined LysC and Trypsin, chymotrypsin only, GluC only, as analyzed using CID-MS2/HCD-MS2 fragmentation. A) Cross-link peptide length by number of amino acids from cross-linked BSA peptide spectra. B) Precursor *m/z* value. C) Number of cross-linked peptide spectra based on charge state. D,E,F,G) Comparison of peptide A and peptide B length from cross-linked peptides identified by trypsin only, combined LysC and trypsin, chymotrypsin only or GluC only protease digestion.

**Table S-1**. List of protein cross-links from in-situ cross-linking of purified chromatin identified from at least 2 out of 3 biological replicates.

